# Palmitoylation of proteolipid protein M6 promotes tricellular junction assembly in epithelia of *Drosophila*

**DOI:** 10.1101/2023.09.04.556077

**Authors:** Raphael Schleutker, Stefan Luschnig

**Affiliations:** Institute of Integrative Cell Biology and Physiology, Cells in Motion (CiM) Interfaculty Centre, University of Münster, D-48149 Münster, Germany

**Keywords:** Epithelium, tricellular junction, cell vertex, palmitoylation, *Drosophila*

## Abstract

Tricellular junctions (TCJs) provide essential adhesive and occluding functions at epithelial cell vertices and play key roles for tissue integrity and physiology, but how TCJs are assembled and maintained is poorly understood. In *Drosophila*, the transmembrane proteins Anakonda (Aka), Gliotactin (Gli), and M6 constitute tricellular occluding junctions. Aka and M6 localize in an interdependent manner to vertices and are required jointly to localize Gli, but how these proteins interact to assemble TCJs was not known. Here, we show that the tetraspan proteolipid protein M6 physically interacts with Aka and with itself. M6 is palmitoylated on a conserved juxta-membrane cysteine cluster. This modification promotes efficient vertex localization of M6 and binding to Aka, but not to itself, and becomes essential when TCJ protein levels are reduced. Abolishing M6 palmitoylation leads to delayed accumulation of M6 and Aka at vertices but does not affect the rate of TCJ growth or mobility of M6 or Aka. Our findings suggest that palmitoylation-dependent recruitment of Aka by M6 promotes initiation of TCJ assembly, while subsequent TCJ growth relies on different mechanisms independent of M6 palmitoylation.

## Introduction

Directional and selective transport across epithelia depends on occluding cell-cell junctions that seal the space between cells to restrict paracellular diffusion. Tight junctions (TJs) in vertebrates and septate junctions (SJs) in invertebrates play analogous roles as diffusion barriers, although they differ in their ultrastructure and molecular composition (reviewed in Higashi and Chiba 2020). Junctions between two adjacent cells (bicellular junctions, BCJs) represent the most abundant intercellular contacts in epithelia. However, where three cells meet at cell vertices, BCJ strands are discontinuous and turn from being oriented parallel to the apical cell surface to align with the central gap, which is flanked by three adjoining cells and sealed by specialized tricellular junctions (TCJs; Staehelin 1973; Fristrom 1982; Graf et al. 1982; Noirot-Timothée et al. 1982). TCJs are composed of a distinct set of proteins that mediate adhesive and occluding properties at cell vertices. In vertebrates, the tetraspan- transmembrane (TM) protein Tricellulin (Ikenouchi et al., 2005) is recruited to tricellular tight junctions (tTJs) by the Angulin family transmembrane proteins Lipolysis-stimulated lipoprotein receptor (Angulin-1/LSR; Masuda et al. 2011) and Immunoglobulin-like domain- containing receptors 1 and 2 (ILDR1/2; Higashi et al. 2013). TCJs play fundamental roles in epithelial barrier function, tissue integrity, cytoskeletal organization, and mitotic spindle orientation (reviewed in Bosveld et al., 2018; Higashi and Chiba, 2020; Higashi and Miller, 2017). Mutations in Tricellulin and Angulin-2/ILDR-1 cause deafness associated with degeneration of cochlear hair cells in mice and humans (Higashi et al., 2013; Riazuddin et al., 2006). Moreover, TCJs provide preferred routes for leukocyte extravasation from blood vessels (Burns et al., 1997; Sumagin and Sarelius, 2010) and are exploited by bacterial pathogens for breaching epithelial barriers (Papatheodorou et al., 2011; Sumitomo et al., 2016). However, despite the key functions of tTJs in epithelial biology and disease, the mechanisms underlying their assembly at cell vertices are only beginning to be understood.

Epithelia of invertebrates also display specialized tricellular occluding junctions. In *Drosophila*, the transmembrane proteins Anakonda (Aka; Byri et al. 2015), Gliotactin (Gli; Schulte, Tepass, and Auld 2003), and M6 (Zappia et al., 2011) organize tricellular septate junctions (tSJs) and are required for epithelial barrier function. Aka and M6 localize to TCJs in a mutually dependent fashion and act jointly to recruit Gli. Gli, in turn, is dispensable for targeting Aka and M6 to vertices, but is required for maintaining Aka and M6 localization, possibly by linking TCJ complexes to adjacent bicellular SJ strands (Esmangart de Bournonville and Le Borgne, 2020; Wittek et al., 2020).

A key open question is which specific features of cell vertices, and which corresponding cellular mechanisms, direct the assembly of TCJ complexes to this small portion of the plasma membrane. In *Drosophila*, Aka mediates homophilic cell adhesion with its triple-repeat-extracellular domain and is required on three adjoining cells for TCJ formation, suggesting that Akás extracellular domain recognizes vertex geometry or the presence of three adjacent plasma membranes (Byri et al., 2015). In addition, negative membrane curvature and a specific lipid composition at vertices may attract specific proteins through interactions mediated by special transmembrane or lipid-binding domains (Aimon et al., 2014). In vertebrates, Angulin-1 is palmitoylated on cytoplasmic cysteine residues, and this post-translational modification is required, along with Angulin-1’s extracellular domain, for targeting to tTJs, possibly by mediating attraction to a cholesterol-enriched membrane domain at vertices (Oda et al., 2020).

S-palmitoylation is the covalent attachment of a C16 acyl chain to cytosolic cysteines of cytoplasmic or integral membrane proteins and is catalyzed by cytoplasmic palmitoyltransferases (reviewed in Chamberlain and Shipston 2015). Unlike other lipid modifications, S-palmitoylation is reversible, and can hence regulate protein distribution or function in a switch-like manner. S-Palmitoylation was shown to regulate partitioning of proteins into specific membrane domains, intracellular trafficking, protein-protein interactions, and activity of TM proteins (reviewed in Blaskovic et al., 2013).

In a search for molecular determinants that promote vertex-localization of TCJ proteins, we found that the *Drosophila* proteolipid protein M6, like its vertebrate homologue GPM6a (Honda et al., 2017), is S-palmitoylated on a conserved cluster of juxtamembrane cysteines, whereas S-palmitoylation is not detectable on other known TCJ components. We show that S-palmitoylation promotes vertex localization of M6 and its direct interaction with Aka. S-palmitoylation of M6 is required for efficient initiation of TCJ assembly, but not for subsequent TCJ growth, thus revealing a distinct role of palmitoylation during an early step of TCJ assembly.

## Results

### Three M6 isoforms are expressed in epithelia and localize to vertices

To identify mechanisms that target M6 to vertices, we analyzed the subcellular distribution of M6 protein isoforms. The *M6* locus encodes six annotated protein isoforms (B through G; Fig. 1A; FlyBase), which differ in their N-terminal cytoplasmic domain. Isoforms B, C, and D were detectable by immunoblot in embryo lysates (Fig. S1A-C) and are expressed in ectodermal (isoforms C and D) and endodermal (isoform B) tissues (Fig. S1D-H).

**Figure 1.**
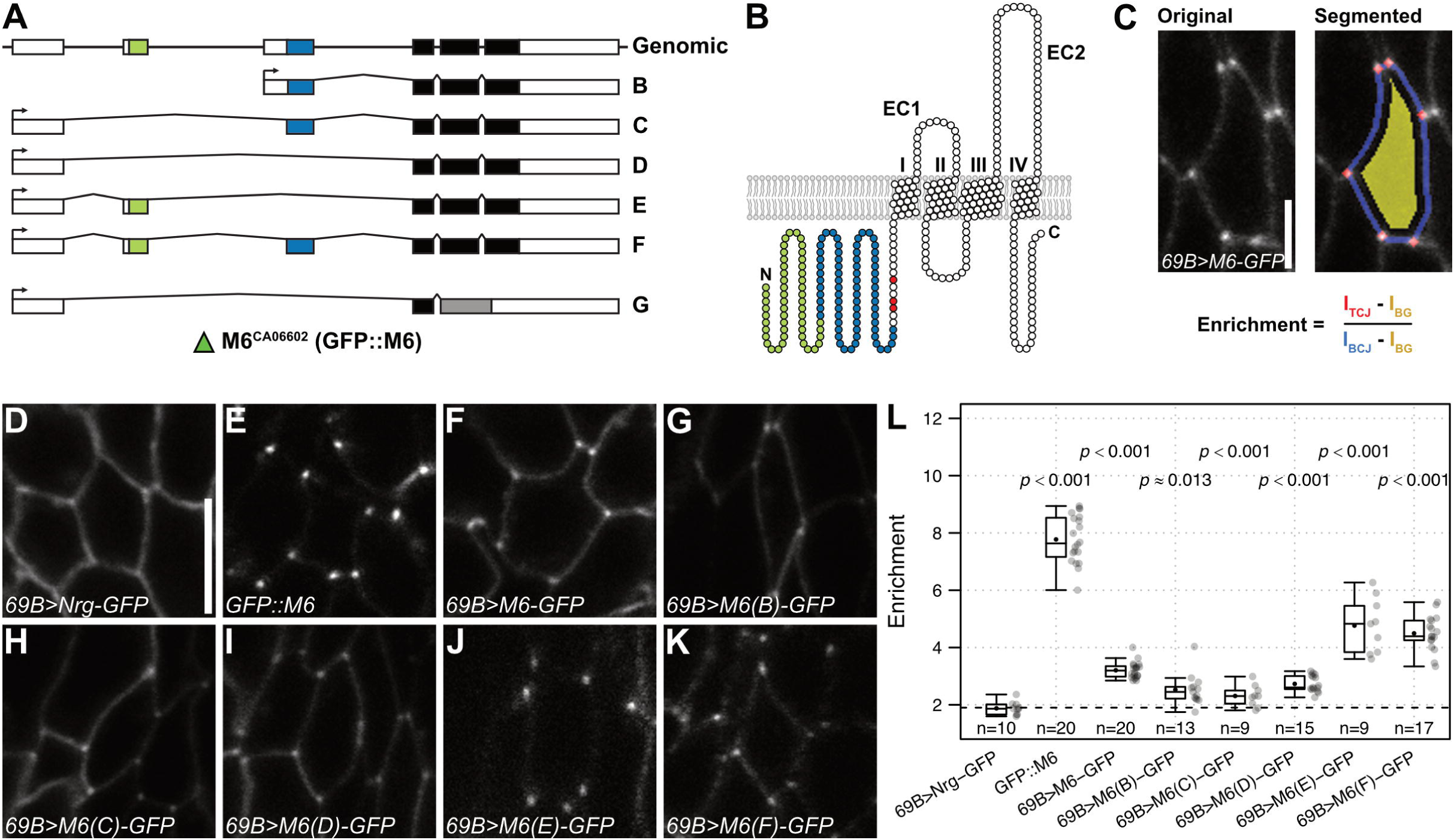
All M6 protein isoforms localize to cell vertices. (**A**) *M6* genomic locus (top) and transcripts encoding isoforms B-G (FlyBase). Exons shared by all isoforms are marked in black, isoform-specific exons are colored. Green triangle marks the M6::GFP (CA06602) protein trap insertion site. (**B**) Topology of M6 protein (isoform F). M6 isoforms comprise four TM domains, extra- and intracellular loops, and a variable N-terminus. Coloring corresponds to exons as in (A). Palmitoylated cysteines are marked in red. **(C)** Analysis of vertex enrichment of M6-GFP fusion proteins. Example of a raw (left) and segmented (right) image of M6-GFP in an epidermal cell is shown. Vertex enrichment is calculated as the ratio of background-subtracted signals at tricellular (TCJ) and bicellular junctions (BCJ). (**D-K**) Distribution of Nrg::GFP (control; D), endogenous GFP::M6 (E), and UAS-M6-GFP (F– K) constructs expressed under control of *69B*-Gal4 in epidermis of living embryos (stage 15). M6-E and M6-F (J and K) show low signals and corresponding images were acquired with higher gain. (**L**) Quantification of vertex enrichment of Nrg-GFP (control), endogenous GFP::M6, and M6- GFP isoforms. Note that vertex enrichment of all M6 isoforms is significantly higher than that of Nrg-GFP. P-values are indicated (pairwise Wilcoxon rank-sum test with Holm correction). Each datapoint represents the mean of one embryo. Number of embryos (n) analyzed is indicated. Box plots here and in following figures show maximum and minimum observations, upper and lower quartile, median (horizontal line), and mean (black dot). Scale bars: (C), 2.5 µm; (D to K), 5 µm.

We tagged each isoform C-terminally with GFP, drove expression of these constructs in the embryonic epidermis (Fig. 1D–K), and determined the enrichment of M6-GFP proteins at vertices as the ratio of signals at tricellular and bicellular contacts (Fig. 1C). This revealed that all M6 isoforms accumulated at vertices with mean enrichment factors of greater than 2.3-fold, whereas the bicellular SJ protein Neuroglian (Nrg-GFP; control) was enriched less than 2-fold (Fig. 1L). Notably, vertex enrichment varied between M6-GFP isoforms and was inversely proportional to overall signal intensity, suggesting that saturation effects upon overexpression impede vertex enrichment. Consistent with this notion, endogenous GFP::M6^CA06602^ showed higher vertex enrichment (7.8-fold; Fig. 1E, L) than the individual overexpressed isoforms. We conclude that all M6 isoforms contain the element(s) required for vertex localization.

### All M6 isoforms support TCJ formation

To test which M6 isoforms support TCJ formation, we expressed each isoform in epidermal stripes in *M6* deficient (*M6^MB02608^/Df(3L)BSC419*) embryos and asked whether the M6 isoforms rescue TCJ localization of Aka and Gli, which are mislocalized along BCJs in the absence of M6 (Fig. S2). cDNAs of *M6* isoforms B, C, and D, as well as an intron-containing UAS-*M6* construct capable of producing all *M6* long-transcript isoforms, rescued TCJ localization of Aka and Gli (Fig. S2A-E). Isoforms M6-E and M6-F showed low expression levels and rescued TCJ localization of Gli, but only partially rescued Aka localization (Fig. S2F,G). However, in principle, all M6 isoforms can support TCJ formation.

### M6 protein is S-palmitoylated on a conserved cluster of cysteine residues

We next asked which elements that are shared by all M6 isoforms mediate vertex localization and TCJ formation. Because S-palmitoylation modulates membrane localization of the vertebrate M6 homologue GPM6a (Honda et al., 2017), we asked whether palmitoylation is involved in vertex localization of *Drosophila* M6. To test whether M6 is palmitoylated, we used an acyl-biotin-exchange (ABE) assay that detects S-palmitoylation of cysteine residues (Wan et al. 2007). After substituting thioester-linked-palmitate for biotin, formerly S-palmitoylated proteins are affinity-purified using Streptavidin beads and are detected by immunoblot (Fig. 2A). To validate this assay, we analyzed extracts from embryos expressing either cytosolic GFP or palmitoylated YFP (palm-YFP) in tracheal cells. Indeed, we detected S-palmitoylation of palm-YFP, but not of GFP (Fig. 2B). We then tested the three known TCJ proteins and found that M6, but not Aka or Gli, is palmitoylated in embryos (Fig. 2C). M6 is predicted to be palmitoylated at a conserved cluster of three cysteines in the N-terminal cytoplasmic portion that is common to all M6 isoforms (Fig. 2D). The corresponding cysteines are palmitoylated in mouse GPM6a (Honda et al., 2017). We mutated the three cysteines to serines in UAS-M6-GFP and tested for palmitoylation of the mutant protein in transfected S2R+ cells. While palmitoylation was readily detectable on wild-type M6-GFP, it was absent from the mutant protein (Fig. 2F), indicating that M6 is palmitoylated on at least one of the three cysteines and that this is the only site of S-palmitoylation in M6.

**Figure 2.**
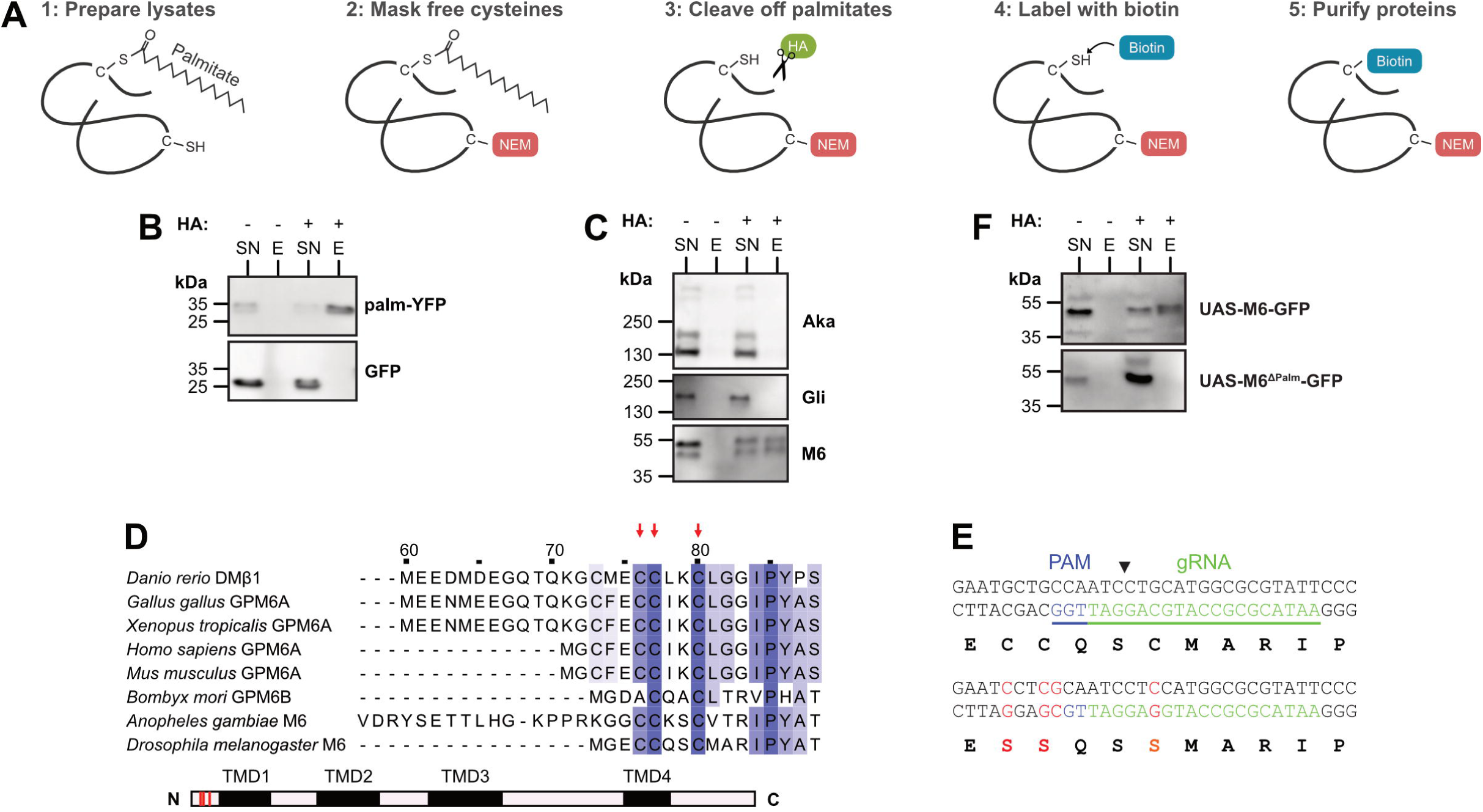
M6 is palmitoylated on a conserved cluster of juxtamembrane cysteine residues. (**A**) Principle of acyl-biotin-exchange (ABE) assay for detecting S-palmitoylation of proteins. Cell or embryo lysates are treated with N-ethylmaleimide (NEM) to mask free thiols, followed by hydroxylamine (HA) treatment to cleave off thioester-linked palmitate, thiol-specific biotinylation, and purification of biotinylated proteins using streptavidin-coupled magnetic beads. (**B**) Lysates of embryos expressing palmitoylated YFP (palm-YFP) or unmodified GFP were subjected to the ABE assay. Eluate (E) and supernatant (SN) fractions after streptavidin purification were analyzed by immunoblot and probed with anti-GFP antibodies. Presence or absence (control) of hydroxylamine (HA) treatment is indicated. Note that S-palmitoylation is detectable on palm-YFP, but not on unmodified GFP. (**C**) ABE assay to test for S-palmitoylation of TCJ proteins Aka, Gli (Gli::YFP), and M6 (GFP::M6) in lysates of embryos (14 to 18 hAEL). Note that M6 (two bands corresponding to isoforms M6-B, M6-C and M6-D), but not Aka or Gli, is S-palmitoylated. (**D**) Multiple sequence alignment of N-terminus of *Drosophila melanogaster* M6 (isoform D) and M6 homologues from indicated species. Note conserved cluster of three cysteines (red arrows) in the juxtamembrane region shared by all *D. melanogaster* M6 isoforms. (**E**) CRISPR/Cas9-based genome editing approach to mutate the three clustered cysteines to serines (red) to generate palmitoylation-deficient M6 mutant (M6^ΔPalm^). Position of guide RNA (underlined), PAM, and cut site (arrowhead) are indicated. (**F**) UAS-M6-GFP or UAS-M6^ΔPalm^-GFP were expressed in S2R+ cells and tested for palmitoylation. Note that M6^ΔPalm^ did not exhibit detectable palmitoylation.

### Palmitoylation promotes accumulation of M6 at vertices

Having established a palmitoylation-deficient M6 mutant (referred to as M6^ΔPalm^), we asked whether palmitoylation is involved in vertex localization of M6. Therefore, we expressed either wild-type M6-GFP or M6^ΔPalm^-GFP in the embryonic epidermis (using *69B*-Gal4) and analyzed the subcellular distribution of the GFP-tagged proteins (Fig. 3A, B). M6^ΔPalm^-GFP was present at the plasma membrane, indicating that palmitoylation is not required for intracellular trafficking or membrane targeting of M6, unlike other proteins, such as Wingless (Wg), which require palmitoylation for their secretion (Franch-Marro et al., 2008). However, M6^ΔPalm^-GFP showed significantly lower (2.2±0.3-fold) vertex enrichment compared to wild-type M6-GFP (3.2±0.3; p<0.001; n=20 embryos; Fig. 3I), suggesting that palmitoylation promotes targeting of M6 to or stabilization at vertices.

**Figure 3.**
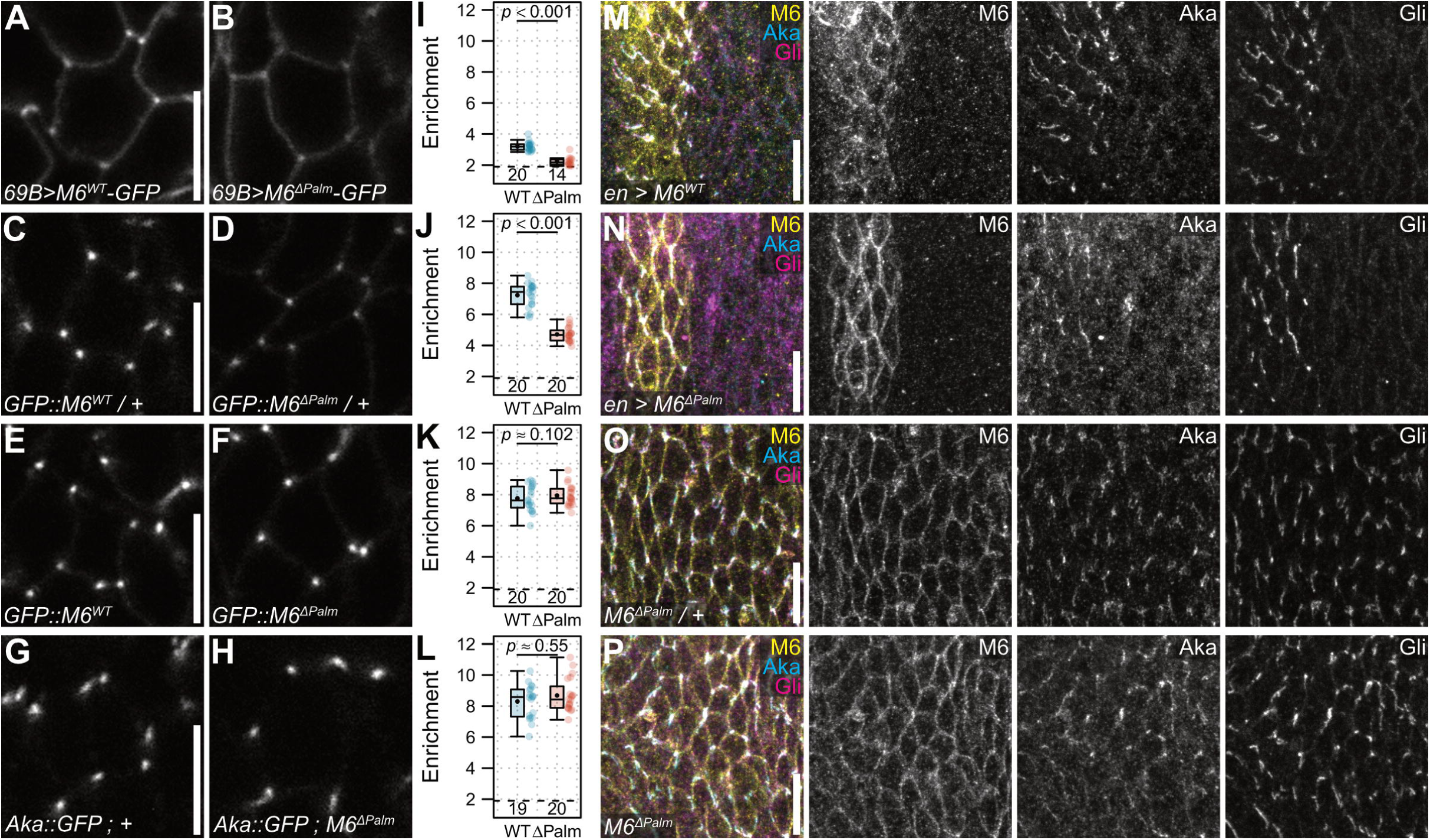
Palmitoylation promotes but is not essential for vertex localization of M6. (**A–H**) *En-face* views of epidermis in living embryos (stage 15) overexpressing wild-type M6- GFP or palmitoylation-deficient M6^ΔPalm^-GFP under control of *69B*-Gal4 (A, B), or expressing endogenous wild-type GFP::M6 (C, E), GFP::M6^ΔPalm^ (D, F), or Aka::GFP (G, H). (**A, B**) When overexpressed in epidermis under control of *69B*-Gal4, M6-GFP (A) shows stronger enrichment at vertices compared to M6^ΔPalm^-GFP (B). (**C, D**) In heterozygous embryos carrying one wild-type copy of untagged M6, endogenous GFP::M6 (C) shows stronger enrichment at vertices compared to GFP::M6^ΔPalm^ (D). (**E, F**) In homozygous embryos lacking untagged M6, endogenous GFP::M6 and GFP::M6^ΔPalm^ show indistinguishable enrichment at vertices. (**G, H**) Vertex enrichment of Aka::GFP in control embryos (G) is indistinguishable from that in *M6*^Δ*Palm*^ homozygous embryos (H). (**I–L**) Quantification of vertex enrichment of GFP signals in the indicated genotypes. Each datapoint represents the mean of 3 to 8 vertices in one embryo. Number of embryos (n) and p-values are indicated. Two-tailed unpaired Student’s *t*-test (J, K) or two-tailed unpaired Wilcoxon rank-sum test (I, L). (**M, N**) *En face* view of lateral epidermis in M6-deficient (*M6^MB02608^/Df(3L)BSC419)* embryos (stage 15) expressing UAS-M6-GFP (M) or UAS- M6^ΔPalm^-GFP (N) in epidermal stripes under control of *en*-Gal4. Embryos were fixed and immunostained against Aka, M6, and Gli. Note that palmitoylation-deficient M6^ΔPalm^ rescues TCJ localization of Gli and partially rescues Aka localization in *en*-Gal4-expressing cells. (**O, P**) *En face* view of lateral epidermis in fixed embryos (stage 15) heterozygous (*M6*^Δ*Palm*^*/+*; O) or homozygous (*M6*^Δ*Palm*^*/M6*^Δ*Palm*^; P) for *M6*^Δ*Palm*^. Embryos were immunostained against Aka, M6, and Gli. Note that TCJ localization of Gli is normal and Aka localization is only mildly affected in *M6*^Δ*Palm*^ homozygotes (P). Scale bars: (A, C, E, and G), 5 µm; (M to P), 10 µm.

To test whether M6 palmitoylation is required for TCJ formation, we expressed M6^ΔPalm^-GFP in epidermal stripes of *M6* deficient embryos and analyzed the distribution of Aka and Gli (Fig. 3M,N). Surprisingly, Aka localization was partially rescued and Gli localization was fully rescued in M6^ΔPalm^-GFP-expressing cells (Fig. 3N), suggesting that M6 palmitoylation is dispensable for TCJ assembly, despite the low enrichment of M6^ΔPalm^-GFP at vertices. However, in this experiment, M6^ΔPalm^ protein was overexpressed, thus impeding conclusions about functional roles of M6 palmitoylation. Therefore, to analyze functions of M6 palmitoylation, we mutated the cysteine cluster in the endogenous *M6* locus (with or without GFP-trap insertion, referred to as *GFP::M6*^Δ*Palm*^ or *M6*^Δ*Palm*^, respectively; Fig. 2E).

### Palmitoylated M6 outcompetes palmitoylation-deficient M6 from localizing to vertices

Endogenous GFP::M6^ΔPalm^ protein showed significantly lower enrichment (4.7±0.5) than wild-type GFP::M6 (7.2±0.8; p<0.001; n=20 embryos) at epidermal cell vertices in heterozygous (*GFP::M6*^Δ*Palm*^*/+*) embryos (Fig. 3C,D,J), resembling the distribution of overexpressed M6^ΔPalm^-GFP (Fig. 3B). Surprisingly, however, in homozygous *GFP::M6*^Δ*Palm*^ embryos, vertex enrichment of GFP::M6^ΔPalm^ was indistinguishable from that of wild-type GFP::M6 protein (Fig. 3E,F,K), suggesting that in the heterozygous situation wild-type M6 protein outcompetes GFP::M6^ΔPalm^ from occupying vertices. Despite this effect, TCJ localization of Aka and Gli appeared normal in *GFP::M6*^Δ*Palm*^ heterozygous and homozygous embryos (Fig. 3O,P), and homozygous *GFP::M6*^Δ*Palm*^ flies were viable and fertile. Thus, although palmitoylation is not essential for vertex localization of M6 or for TCJ formation, it promotes transport to or maintenance of M6 at vertices.

### TCJ assembly is delayed in the absence of M6 palmitoylation

To test this hypothesis, we analyzed the dynamics of GFP::M6 accumulation at epidermal cell vertices in time-lapse movies (Fig. 4A). At the onset of TCJ formation, GFP::M6 accumulates at the apical tip of each vertex in a single spot that subsequently extends basally with a speed of 0.09 µm/min (n=15 vertices in 3 embryos; Fig. 4A’, D; Video S1). This process starts at a small number of vertices during stage 13 and subsequently sweeps across the epidermis over the course of 95±10 minutes (n=3 embryos) until all vertices are occupied by M6 during stage 14 (Fig. 4A,C; Video S1; Wittek et al. 2020). To quantify the kinetics of M6 vertex localization, we determined in each movie frame the fraction of vertices marked by accumulations of GFP::M6 (Fig. 4A). While wild-type GFP::M6 occupied all vertices at 95±10 minutes (n=3 embryos) after the onset of TCJ formation, GFP::M6^ΔPalm^ reached complete vertex occupancy with a delay of 76% after 167±21 min (delay of 26 minutes in half-maximal occupancy time, p<0.05; Fig. 4B, C), indicating that effcient TCJ formation depends on M6 palmitoylation.

**Figure 4.**
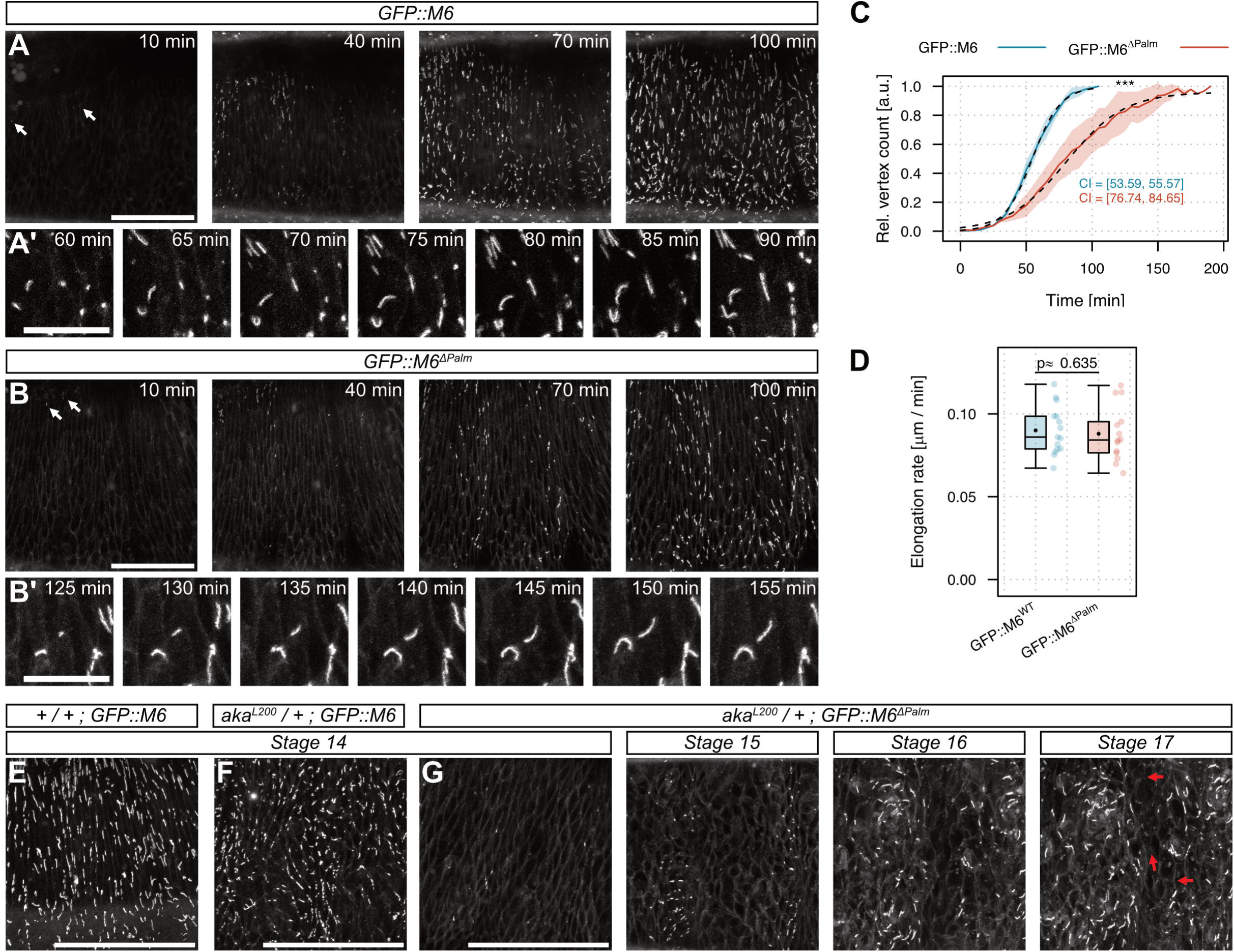
Lack of M6 palmitoylation leads to delayed initiation of TCJ assembly but does not affect the rate of TCJ growth. (**A, B**) *En face* view of dorso-lateral epidermis in living embryos (stage 13) homozygous for either *GFP::M6* (A) or *GFP::M6*^Δ*Palm*^ (B). White arrows denote first appearance of GFP::M6 accumulation at vertices. (**A’, B’**) Close-up of vertices showing extension of GFP::M6 (A’) or *GFP::M6^ΔPalm^* (B’) signals along the apicobasal axis. Time is indicated. Rate of extension was quantified in (D). (**C**) Quantification of vertex accumulation of GFP::M6 (blue) and GFP::M6^ΔPalm^ (red). The number of GFP-positive vertices was counted at each timepoint and normalized to the maximum number of GFP-positive vertices at the end of each movie. Note that GFP::M6 accumulated at most vertices throughout the epidermis within 100 min, whereas vertex accumulation of GFP::M6^ΔPalm^ was delayed. Dashed lines indicate fitted logistic model. Confidence intervals (CI; upper and lower limit) for time of half-maximal accumulation are indicated. n=3 embryos per genotype. (**D**) Quantification of extension rate of GFP::M6 (blue) and GFP::M6^ΔPalm^ (red) along vertices. Note that GFP::M6 and GFP::M6^ΔPalm^ extend with similar rates. n=15 vertices from 3 embryos per genotype. One-tailed *t*-test. (**E,F,G**) *En face* view of living embryos (stage 14). GFP::M6 occupies all vertices by stage 14 in wild-type (E) and *aka^L200^* heterozygotes (F), whereas GFP::M6^ΔPalm^ accumulates only at few vertices at stage 14 in *aka^L200^/+* heterozygotes (G). Note that many vertices fail to accumulate (red arrows) GFP::M6^ΔPalm^ even in late (stage 17) embryos. Scale bars: (A, B, E, F, and G), 50 µm; (A’ and B’), 2 µm.

### M6 palmitoylation promotes initial accumulation, but not extension of M6 clusters at vertices

Interestingly, although the rate of vertex occupancy by GFP::M6 was impaired by the lack of M6 palmitoylation, there was no detectable effect on the rate of extension along a vertex after GFP::M6 has accumulated at its apical tip (0.088 µm/min; n=15 vertices from 3 embryos; Fig. 4D). This suggests that efficient initial accumulation of M6 at vertices depends on palmitoylation, while the subsequent extension of M6 clusters during TCJ growth is independent of M6 palmitoylation. Consistent with this notion, the lack of M6 palmitoylation had only a slight, if any, effect on the mobility of M6 (GFP::M6^ΔPalm^) or of Aka (Aka::GFP) proteins at vertices, as measured by Fluorescence Recovery After Photobleaching (FRAP) experiments (Fig. S3), whereas complete loss of M6 protein was previously shown to lead to substantially increased mobility of Aka and abolishes TCJ formation (Wittek et al., 2020).

### M6 palmitoylation becomes essential when dosage of Aka is reduced

Because vertex accumulation of M6 depends on Aka (Wittek et al., 2020), we wondered whether the delayed TCJ assembly in the absence of M6 palmitoylation can be further perturbed by reducing the dosage of Aka. Indeed, removing one copy of *aka* in the absence of M6 palmitoylation aggravated the delay in vertex occupancy of *GFP::M6*^Δ*Palm*^, with many vertices failing to accumulate M6 even in late-stage embryos (Fig. 4G), and led to synthetic embryonic lethality, indicating that M6 palmitoylation becomes essential when TCJ protein concentration is reduced. Taken together, these results highlight the requirement of M6 palmitoylation for robust TCJ assembly, and raised the question whether Aka and M6 proteins interact with each other, possibly in a palmitoylation-dependent manner.

### M6 interacts with Aka and with itself

To address this question, we carried out co-immunoprecipitation (co-IP) experiments using GFP-tagged M6 as a bait protein (Fig. 5A). We found that GFP::M6 specifically co-precipitated Aka from embryo extracts (Fig. 5B), suggesting that Aka and M6 are part of a protein complex. This interaction was detected also in transfected S2R+ cells expressing UAS-Aka (prey) and UAS-M6-GFP (bait) but no other known junction components, suggesting that Aka and M6 interact directly (Fig. 5D,G). We further noticed that GFP::M6 specifically co-precipitated also the untagged M6-B isoform expressed in embryos (Fig. 5B, arrow). Likewise, M6-GFP co-precipitated 2X-hemagglutinin-tagged M6 (M6-2XHA) in extracts from transfected S2R+ cells (Fig. 5E), indicating that M6, like vertebrate GPM6a (Formoso et al., 2015), interacts with itself.

**Figure 5.**
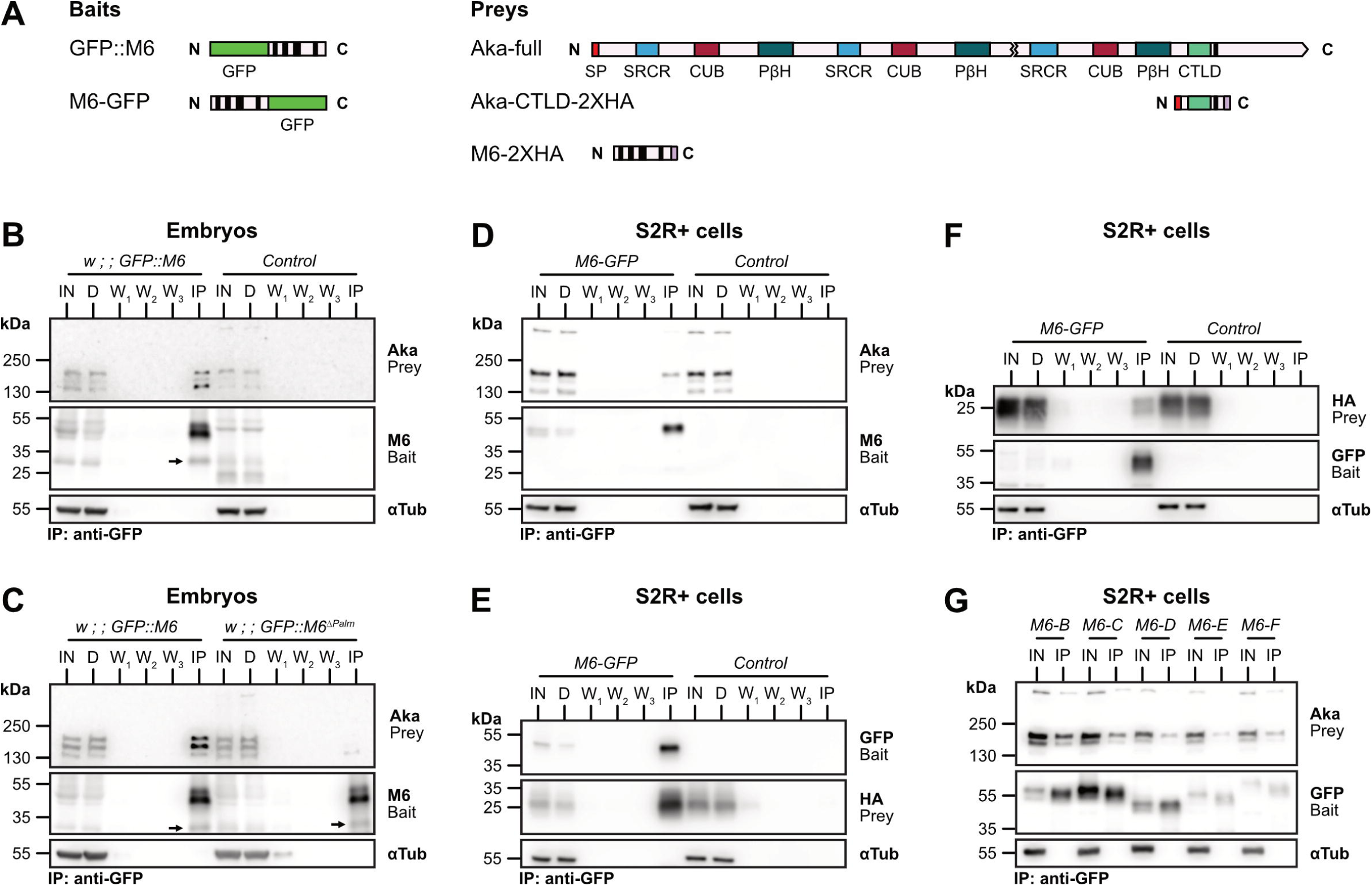
M6 interacts with Anakonda and with itself. **(A)** Schemes of proteins used as bait or prey in co-immunoprecipitation (co-IP) experiments. Akás extracellular portion comprises an N-terminal signal peptide (SP), Scavenger receptor- cysteine-rich (SRCR) domains, complement C1r/C1s, UEGF, BMP1 (CUB) domains, right- handed parallel β-helices (PβH), and a C-type lectin-like domain (CTLD). Black bars indicate transmembrane domains. (**B–G**) IP experiments with lysates from embryos (B,C) or transfected S2R+ cells (D–G) expressing the indicated bait proteins. Immunoblots of input (IN), depleted input (D), washes (W), and immunoprecipitation (IP) fractions were probed with antibodies indicated to the right. Alpha-tubulin was used as a loading control. Sizes (kDa) of a molecular weight marker are indicated to the left. **(B)** IP with anti-GFP antibodies to immunoprecipitate GFP::M6. Note that GFP::M6 co- precipitates Aka and the non-GFP-tagged M6-B isoform (arrow) from lysates of embryos expressing GFP::M6 but not from negative control (*y w*) embryos. **(C)** IP of wild-type GFP::M6 (left) or palmitoylation-deficient GFP::M6^ΔPalm^ (right). Note that co-precipitation of Aka is strongly reduced with GFP::M6^ΔPalm^ compared to GFP::M6, whereas co-precipitation of M6-B (black arrow) is unchanged, indicating that the Aka-M6 interaction is palmitoylation-dependent, whereas M6-M6 interactions are palmitoylation-independent. **(D)** IP experiment with S2R+ cells co-transfected either with M6-GFP (bait) and Aka (prey; left) or with Aka alone (control; right). Note that M6-GFP specifically co-precipitates Aka. **(E)** IP experiment with S2R+ cells co-transfected either with M6(D)-GFP (bait) and M6(D)- 2XHA (prey; left) or with M6(D)-2XHA alone (control; right). Note that M6(D)-GFP specifically co-precipitates M6(D)-2XHA, indicating that M6 self-interacts. (**F**) A short fragment of Aka (scheme in A) comprising the extracellular C-type lectin-like domain (CTLD), the TM domain, and a cytosolic 2XHA tag was co-transfected with UAS-M6- GFP into S2R+ cells. M6 specifically co-precipitates the short Aka fragment. (**G**) Each M6 isoform (B, C, D, E, F) was tagged with GFP (bait) and was individually co-transfected with Aka (prey) into S2R+ cells. Note that all M6 isoforms co-precipitate Aka.

### M6 palmitoylation is required for M6-Aka interaction, but not for M6-M6 interaction

To test whether palmitoylation of M6 influences its binding to Aka or to itself, we performed co-IP experiments with extracts from *GFP::M6*^Δ*Palm*^ embryos (Fig. 5C) or from *M6*^Δ*Palm*^-GFP- transfected S2R+ cells (Fig. 5D). Interestingly, the Aka-M6 interaction was strongly reduced or absent, whereas the M6-M6 homotypic interaction was unaffected (Fig. 5C, arrow), suggesting that M6 interacts with Aka in a palmitoylation-dependent manner, but self-interacts in a palmitoylation-independent manner. Aiming to narrow down the M6-interacting site in Aka, we found that an Aka fragment comprising only the C-type-lectin domain (CLTD) and the TM domain (Aka-CTLD-2XHA) but lacking the remaining extracellular and cytosolic parts of Aka (Fig. 5A), still co-precipitated with M6-GFP (Fig. 5F). This suggests that Aka and M6 interact either via their TM domains or via Aka’s CTLD and implies that the palmitoylated cytosolic cysteine-cluster in M6 is not the site of interaction with Aka. Instead, M6 palmitoylation appears to influence the interaction with Aka in an allosteric fashion.

## Discussion

How specialized junctional complexes with distinct adhesive and occluding properties are built at three-cell-contacts remains a fundamental open question in epithelial biology (Bosveld et al., 2018; Higashi and Chiba, 2020; Higashi and Miller, 2017). Three TCJ-specific proteins, Aka, Gli and M6, are known in *Drosophila*, but how they interact to assemble TCJs was not known. Here we report insights into the early steps of TCJ assembly, which depends on interactions between the transmembrane proteins Aka and M6. We show first that all M6 isoforms share the elements required for vertex localization and TCJ formation. Second, we demonstrate that M6 is palmitoylated on a conserved cysteine cluster and that this modification is required for efficient initial localization of M6 to vertices, but not for subsequent TCJ growth and maintenance, indicating that different mechanisms control the early and late phases, respectively, of TCJ formation. Third, we show that Aka interacts with M6 in a palmitoylation-dependent manner, while M6 interacts with itself independent of palmitoylation, possibly forming homotypic clusters like the mammalian M6 homologue GPM6a.

These findings provide a molecular basis for the interdependent localization of Aka and M6 at TCJs, although it is not yet clear which physical or chemical features of vertices mediate localization of proteins to these sites. Aka engages in homotypic trans-interactions that depend on its expression in three adjacent cells and on Akás large extracellular domain, but not on its cytoplasmic domain for accumulating at vertices (Byri et al., 2015). This suggests that the specific geometry of vertices or the presence of three adjacent plasma membranes act to recruit or maintain Aka. M6 is required for vertex localization of Aka in a permissive fashion (Wittek et al., 2020). Our findings suggest possible mechanisms underlying this role. In one scenario, M6 might stabilize Aka in the plasma membrane by preventing its internalization. Supporting this idea, in the absence of M6 Aka was barely detectable at the plasma membrane but instead distributed throughout the cytoplasm (Wittek et al. 2020). Alternatively, association with M6 might concentrate Aka in a vertex-specific plasma membrane domain and thereby facilitate *trans*-interactions between Aka molecules on adjacent cells. This is reminiscent of the role of tetraspanins, which interact laterally among themselves and with other proteins to form large assemblies called tetraspanin-enriched microdomains (TEMs; Hemler 2003) or tetraspanin web (Lagaudrière-Gesbert et al., 1997) that control the clustering and activity of TM proteins, such as EGFR or integrins (reviewed in van Deventer et al., 2017).

We identified S-palmitoylation as a post-translational modification, which, although not essential for vertex localization of M6 or for TCJ formation, significantly enhances the efficiency and robustness of this process. Supporting this notion, palmitoylation of M6 becomes essential for TCJ formation and viability under conditions of reduced TCJ protein concentration. How could palmitoylation promote targeting of M6 to vertices? In many TM proteins, including claudins (Van Itallie et al., 2005), tetraspanins (Yang et al., 2004), and mammalian GPM6a (Honda et al., 2017), palmitoylation of juxtamembrane cysteines mediates association with cholesterol-rich membrane domains (Blaskovic et al., 2013). Negative plasma membrane curvature at vertices might favor accumulation of cholesterol (Yesylevskyy et al., 2017) and could thereby attract palmitoylated proteins. Consistent with this idea, methyl-beta-cyclodextrin-induced depletion of cholesterol from the plasma membrane impaired the vertex localization of palmitoylated Angulin-1 in cultured mammalian cells (Oda et al., 2020), although direct evidence for elevated cholesterol levels at cell vertices is lacking thus far.

Lack of M6 palmitoylation reduces, although it does not completely abolish, the interaction with Aka, resulting in delayed accumulation of M6 and Aka at vertices. How could palmitoylation affect the interaction of M6 with Aka? Structural predictions using AlphaFold (Jumper et al., 2021) and PPM3 (Predicting Protein position in Membranes; Lomize et al., 2022) suggest that M6 protein adopts a tilted orientation in the membrane (Fig. S4). Palmitoylation of the juxtamembrane cysteine cluster could stabilize the tilt and help to resolve a potential hydrophobic mismatch resulting from the different lengths of the four TMDs, thereby promoting a confirmation of M6 that is favorable for interaction with Aka. Palmitoylation of LRP6 was proposed to act in this way by resolving a hydrophobic mismatch that otherwise causes the protein to be retained in the ER (Abrami et al., 2008). Similarly, tetraspanins, *e.g*., CD151, require palmitoylation for interaction with other transmembrane proteins (Zevian et al., 2011).

Intriguingly, eliminating palmitoylation of M6 revealed a distinct requirement of this modification for efficient initial accumulation of M6 at vertex tips, whereas the subsequent extension of these clusters along the apical-basal vertex axis apparently relies on a different mechanism. This may include homotypic M6-M6 interactions, which we found to be independent of M6 palmitoylation and which may aid in organizing an M6-enriched plasma membrane domain, reminiscent of TEMs (Hemler 2003). Interestingly, in mammalian cells, palmitoylation of juxtamembrane cysteines was proposed to play an analogous role in targeting Angulin-1 to cholesterol-enriched membrane domains at vertices, where Angulin-1 recruits Tricellulin (Oda et al., 2020). However, Angulin-1 vertex localization appears to be dispensable for maintaining Tricellulin localization after its initial recruitment to TCJs (Oda et al., 2020), suggesting that Tricellulin is maintained at vertices through association with other factors, *e.g*., with Claudin-based tight-junction strands (Ikenouchi et al., 2008). Analogously, in *Drosophila*, M6 palmitoylation is required early during TCJ formation for efficient targeting of M6 and Aka to vertices, whereas Aka is subsequently stabilized at TCJs through interaction with components of septate junction limiting strands, including Gli (Esmangart de Bournonville and Le Borgne, 2020). Thus, although TCJs of vertebrates and invertebrates comprise different sets of proteins, our findings suggest that in both cases juxtamembrane palmitoylation contributes to targeting TM proteins to cell vertices during TCJ assembly.

## Materials and Methods

### Drosophila husbandry

*Drosophila* was maintained on cornmeal-agar medium with added dry yeast. Embryos were collected on apple juice agar plates at 22°C or 25°C. The sex of embryos was not assessed.

### *Drosophila* strains and genetics

*Drosophila* stocks are described in FlyBase, unless noted otherwise: *aka^L200^* (Byri et al., 2015), *M6^CA06602^* (Buszczak et al., 2007), *M6^MB02608^* (Bellen et al., 2011), Aka::GFP, UAS-M6, UAS-M6-GFP (Wittek et al., 2020), UAS-Nrg^167^-GFP (Byri et al., 2015), UAS-GFP, UAS-palm-YFP, *UAS-mCherry-nls, 69B-Gal4*, *btl*-Gal4, *en-Gal4, Df(3L)BSC419*, *CyO Dfd-GMR-nvYFP, TM6b Dfd-GMR-nvYFP* (Le et al., 2006), *TM2 Delta2-3*, *nos-Cas9*.

### Genome editing

A CRISPR-Cas9-based strategy was used to mutate the three clustered cysteines to serines in the *M6* locus of the *GFP::M6^CA06602^*line to yield *GFP::M6*^Δ*Palm*^. The sgRNA sequence 5’- GAATACGCGCCATGCAGGAT was cloned in the pCFD3 vector (Port et al., 2014). The resulting sgRNA plasmid (300 ng/µl) along with the ssDNA repair template (5’- ATTGCATTCAAAAGTTTATTGATATTATTTTTCCTAAAAATGTTTTTAGGAGAAT<u>c</u>CT<u>cg</u>CAA TCCT<u>c</u>CATGGCGCGTATTCCCTACGCCACCCTGATAGCCACTCTGATGTGTCTCCTG; mutations underlined, 100 ng/µl, Eurofins Genomics) was injected into *nos-Cas9;; GFP::M6^CA06602^* embryos. Individual F_2_ males were screened for successful conversion by amplifying from genomic DNA of single adult flies a 795bp fragment using a forward primer (5’-GAGAAT<u>c</u>CT<u>cg</u>CAATCCT<u>c</u>CAT) that is specific to the mutated sequence and a reverse primer (5’-GCTCCTGGAACTTTTCGTGG). Positive candidates were analyzed by sequencing the target region. One line containing all four mutations was kept. In this line, the *w^+^*-marked GFP-trap P-element (P{PTT-GA}; Buszczak et al. 2007) was excised by P-element transposase to generate non-GFP-tagged *M6*^Δ*Palm*^.

### Molecular biology

Coding sequences for each annotated *M6* isoform (FlyBase) were synthesized (GenScript) with a 5’ EcoRI site and 3’ (GGS)_5_ linker and NotI site. *M6* isoforms were fused either with the GFP coding sequence (flanked by 5’ NotI and 3’ XbaI sites) or with a 2XHA tag sequence (flanked by 5’ NotI and 3’ XbaI sites) and subcloned into EcoRI/XbaI digested pUASt-attB vector (Bischof et al., 2007), such that each M6 isoform was tagged C-terminally either with GFP or with 2XHA.

Point mutations to change cysteines to serines were inserted into the pUASt-attB-M6-GFP plasmid (Wittek et al., 2020) using site-directed mutagenesis. The plasmid was amplified using mutation-bearing oligonucleotides (forward primer: 5’-GAGAAT<u>c</u>CT<u>c</u>CCAATCCT<u>c</u>CATGGCGC; complementary reverse primer). Bacterial template DNA was digested with DpnI.

All UAS constructs were inserted into the attP2 genomic landing sites using PhiC31- mediated site-specific integration (Bischof et al., 2007).

### Cell culture

*Drosophila* S2R+ cells were cultured in Shields and Sang M3 medium (Biomol) supplemented with Penicillin-Streptomycin and fetal calf serum. 1×10^6^ cells were seeded per well in 24-well plates (Sarstedt) one day before transfection. Cells were transfected with act5c-Gal4 plasmid (600 ng) and pUAST-based expression plasmid (1000 ng) using FugeneHD reagent (Promega). Transfected cells were cultured at 25°C and processed within 1 to 3 days.

### Immunostaining

Embryos were dechorionated in sodium hypochlorite, washed, and fixed in 4% formaldehyde in PBS/heptane for 20 min at room temperature and then devitellinized by shaking in methanol/heptane. The following primary antibodies were used: guinea pig anti-M6 serum #1 (1:3000; Wittek et al. 2020), guinea pig anti-M6 serum #2 (1:3000; this work), rabbit anti-Aka (1:250; Byri et al., 2015), mouse anti-Gli IF6.3 (1:500; Schulte et al. 2003). Goat secondary antibodies were conjugated with Alexa 488, Alexa 568, or Alexa 647 (1:500; ThermoFisher). GFP signal was enhanced with FluoTag-X4 anti-GFP (1:500; NanoTag).

### Antisera

The anti-M6 antiserum #2 was generated by immunizing guinea pigs (Eurogentec) with the peptides GKGNNRDRIRDPRE and RRNSYRSDHSLDRYT (corresponding to aa 57–71 and 102–116 in M6 isoform F) conjugated to keyhole limpet hemocyanin. Final serum was used at a dilution of 1:3000 for immunostainings.

### Acyl-biotin-exchange assay

The acyl-biotin-exchange (ABE) assay was adapted from (Wan et al., 2007). Approximately 200 embryos (16-20 hAEL) were collected, dechorionated, and homogenized using a 7.5 ml Dounce homogenizer in 1.2 ml lysis buffer containing 1.7% Triton X-100, 10 mM N-ethylmaleimide (NEM) and 1x protease inhibitor (ThermoFisher). GFP-tagged M6 was purified using 20 µl GFP-Trap magnetic agarose (Chromotek). For other proteins, lysates were further processed without an intermediate purification step. Further steps including hydroxylamine (HA) treatment and biotinylation were carried out as described in (Wan et al., 2007). Biotinylated proteins were purified using 10 µl streptavidin-coupled magnetic beads (ThermoFisher) and eluted in 40 µl 1x Lämmli buffer at 70°C for 10min. Samples were applied to SDS-PAGE followed by Western blotting and staining for detection of palmitoylated proteins.

### Co-immunoprecipitation

Co-immunoprecipitation experiments were performed with extracts from embryos or from transfected S2R+ cells. Approximately 500 embryos (14 to 18 hAEL) were dechorionated and homogenized using a 7.5 ml Dounce homogenizer in 600 µl lysis buffer (1% Brij-97, 0.1% Tween-20, 10 mM Tris/HCl, 150 mM NaCl, 1 mM MgCl_2_, 1 mM CaCl_2_, 1x protease inhibitor; pH 7.4) on ice. S2R+ cells were lysed in 200 µl lysis buffer and otherwise treated like embryo extracts. Lysates were incubated at 4°C with end-over-end rotation for 30 min followed by centrifugation (2000xg, 5 min, 4°C). Supernatants were incubated for 30 min with 20 µl GFP-Trap magnetic beads (Chromotek gtd-20) at 4°C with end-over-end rotation. Beads were washed three times with 200 µl lysis buffer. During the last washing step, beads were transferred to a new vial and NaCl concentration was raised to 300 mM. Proteins were eluted in 60µl 1x Lämmli buffer by incubating for 10 min at 70°C.

### Immunoblots

Samples were mixed 3:1 with 4x Lämmli buffer and incubated at 70°C for 10 min. 14µl were loaded on 4%-20% Mini-PROTEAN TGX gels (Bio-Rad) and blotted on 0.45µm PVDF membranes (Amersham). The following primary antibodies were used: mouse anti-GFP JL-8 (1:1000; TaKaRa), mouse anti-Tubulin AA4.3-c (1:5000; DSHB), rabbit anti-Aka (1:1000; Byri et al., 2015), guinea pig anti-M6 serum #1 (1:2500; Wittek et al., 2020). Primary antibodies were detected with HRP-conjugated secondary antibodies (1:10000; Thermo Fisher) and ECL Prime kit (Amersham). Raw images were adjusted for brightness and contrast using ImageJ (Schindelin et al., 2012).

### Microscopy

Imaging was performed on a Leica SP8 confocal microscope with a 40x/1.3 NA oil immersion objective. For live imaging, dechorionated embryos were staged according to gut morphology, mounted on glue-coated coverslips (0.17mm, grade #1.5), and covered with Voltalef 10S oil (VWR).

### Image analysis

Images were processed using either FIJI (Schindelin et al., 2012) or Python 3.9 with the Scikit-Image package v0.18.2 (Walt et al., 2014). Image panels in figures were assembled using OMERO.figure v4.4.1 (Allan et al., 2012).

### Quantfication of vertex enrichment

Enrichment of fluorescently tagged proteins at cell vertices was determined as follows. Embryos (stage 15) were aligned laterally and mounted as described above. Z-stacks (15 slices with 768x768px each) through epidermal cells were acquired at 100Hz. In each stack several cells were analyzed using a semiautomatic pipeline. Vertices were marked manually in the slice with highest vertex intensity and segmented by selecting a circular area of 5px diameter around the marked point. Using the marked vertices as starting points the bicellular membrane was then segmented automatically using a self-written Python script. For each cell, the mean intensity at vertices (*I_TCJ_*), the bicellular membrane (*I_BCJ_*), and in the cytoplasm (*I_BG_*) was measured (see Fig. 1C). Enrichment was calculated as

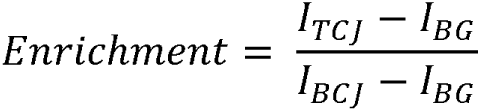

Per embryo, three to eight cells were selected and the mean for each embryo was calculated. Only vertices bordered by three cells but not four or more cells were considered. Sample size (n) indicated in the figures states the number of analyzed embryos.

### Quantification of TCJ initiation and junction growth

Staged embryos were mounted dorso-laterally. The thorax region of embryos was imaged using a 40x/1.3 NA oil immersion objective and 2.5x zoom, 200 lines/sec speed, 1528x1528px resolution, 18 slices with a step-size of 0.5µm, at 5min intervals. Movies were processed frame by frame by applying a Gaussian filter (standard deviation 1.5) followed by a white top-hat transform with a circular kernel of 9px diameter. Movies were afterwards binarized using Otsu’s threshold based on the stack at the last timepoint. Objects smaller than 10px were filtered out. Afterwards, we counted the number of objects at each timepoint and normalized the count to the highest number of objects found at any timepoint yielding the relative vertex count. All movies were aligned temporally such that the first timepoint with a relative vertex count of at least 0.05 was set to 30min. This data was used to fit a logistic regression model using the *nls* and *SSlogis* function of the R statistical software package (v4.1). A 95% confidence interval was calculated for the *xmid* parameter, which represents the timepoint with a relative vertex count of 0.5. While an overlap of the intervals does not allow statements about statistical significance, a non-overlap indicates statistically significant differences in accumulation time to a test-niveau of α = 0.05. Extension of GFP signals along the apico-basal vertex axis was analyzed at manually selected vertices that were oriented mostly parallel to the XY-plane and grew throughout the entire observation phase. TCJ length was measured in 3D shortly after initial accumulation and 30min later by manually tracing GFP::M6 signal through Z stacks. TCJ growth rate was calculated by dividing the measured length increase by the observation time.

### Fluorescence recovery after photobleaching

Stage 15 embryos were selected according to gut morphology and were mounted dorsally. Movies were acquired using a 40x/1.3 NA oil immersion objective using the FRAP wizard of the LAS X software (Leica) with a speed of 400 lines/s and 2x line accumulation. For each embryo, three pre-bleach stacks (each with four z-slices) were acquired followed by three successive bleach rounds at 100% laser intensity followed by 41 post-bleach stacks. The interval between pre- and post-bleach stacks was 30s. Z-shift was corrected manually during acquisition. Since the FRAP data of vertices is not suitable for classical FRAP analysis we used a simplified approach to model the recovery. For each timepoint *t* we generated average intensity projections of slices 2-4. The vertex of interest was selected manually in each frame. We measured the intensities using circular regions of interest with a radius of 5 pixels. The FRAP values were calculated as follows. Let be *I_t_*, *t* ∈ {-3,-2,…,40}, the raw intensities at timepoint t (bleached between *t* = -1 and t = 0). Let be *I*_0.95,t_ the 95% intensity quantile of the frame (to account for bleaching during acquisition) and *b_t_* the background intensity at timepoint t. Then the corrected intensities *C*_t_ were calculated as

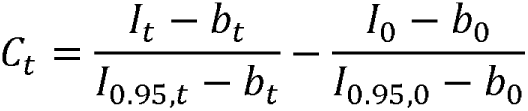

The normalized intensities *N_t_* were calculated as

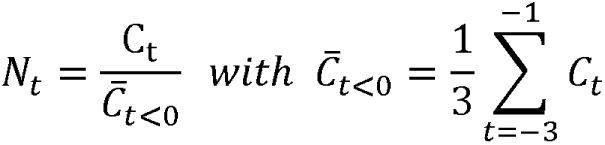

The graphs show the mean values of *N_t_*± *SD_t_*. The recovery function *f*_A,τ_ is defined as

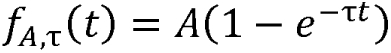

Here,*A* = represents the mobile fraction and τ defines the fluorescence recovery rate. *A* = and τ were fitted numerically with a non-linear least square approach using the *nls* function of the R statistical software package (v4.1). The mobile fraction was directly inferred from =. The half-recovery time was calculated as 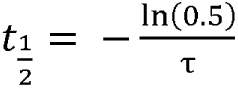.

### Statistics

Sample groups were checked for normality using the Shapiro-Wilk test and for equal variances using an F test. In case of normality and equal variances a pairwise two- or one-sided *t*-test was applied. In case of unequal variances, a *t*-test using pooled standard deviations was used (Welch-test). If data was not normally distributed, a Wilcoxon rank-sum test was applied. If samples were used for several tests, a p-value correction according to Holm (Holm, 1979) was applied to correct for multiple testing.

## Author contributions

Conceptualization, all authors; Methodology, all authors; Investigation, R.S.; Formal Analysis, R.S.; Visualization, R.S.; Writing – Original Draft, R.S.; Writing – Review and Editing, S.L.; Funding Acquisition, S.L.; Supervision, S.L.

## Acknowledgements

We thank Wilko Backer for expert technical help and Vanessa Auld, Manuel Hollmann, Anne Uv, and Tian Xu for providing antibodies and fly stocks.

## Data and materials availability

Fly stocks, reagents, and custom Python scripts described in this study are available upon request.

## Supplemental Figures

**Figure S1 (related to Figure 1) Expression of M6 isoforms in embryos.**

**(A)** Scheme of *M6* transcript isoforms. Insertion site of the GFP::M6 (M6^CA06602^) protein trap transposon and positions of peptides used to generate anti-M6 serum #1 and anti-M6 serum #2 are indicated by triangles. The GFP::M6 protein trap tags tags isoforms from the upstream promoter (all but isoform B). Predicted isoform G contains a retained intron and downstream premature stop codon, resulting in a predicted truncated protein, and was not further considered.

**(B)** Immunoblot of extracts from embryos (*y w*; 0-24h after egg lay) or from S2R+ cells transfected with single HA-tagged M6 isoforms indicated on top. Two bands (green arrows) in the embryo lysate correspond to isoforms B or C (36 kDa) and D (28 kDa), respectively (red arrows).

**(C)** Immunoblot of extracts from embryos (0-24h after egg lay) expressing endogenous GFP::M6 or from S2R+ cells transfected with the indicated GFP-tagged M6 isoform. Three bands (green arrows) correspond to untagged isoform B (lower band; compare panel A), tagged isoform D (middle band), and tagged isoform C (upper band).

**(D)** Embryo (stage 16) expressing GFP::M6 (M6^CA06602^) immunostained against GFP. GFP::M6 tags M6 isoforms C, D, E, F, and G and is detected in ectodermal tissues, including epidermis (ED), salivary glands (SG) and tracheal dorsal trunk (DT), but not in endodermal midgut (MG).

**(E, F)** Embryos (stage 16) heterozygous (E) or homozygous (F) for *Df(3L)BSC419.* Immunostaining with anti-M6 serum #1 detects M6 expression in the ectoderm (epidermis, salivary gland, trachea) resembling distribution of the GFP::M6 protein trap (D), and additionally in the endoderm (midgut). Absence of signal in *Df(3L)BSC419* homozygous embryo (F) indicates specificity of antiserum. Asterisk in (E) indicates YFP signal from the *Dfd-GMR-YFP* balancer chromosome used to genotype embryos.

**(G, H)** Embryos (stage 16) heterozygous (G) or homozygous (H) for *Df(3L)BSC419.* Immunostaining with anti-M6 serum #2 detects M6 expression in endoderm (midgut), but not in ectodermal tissues. Note that endodermal expression corresponds to M6-B, which is the only isoform not tagged by the GFP::M6 (M6^CA06602^) protein trap. Signal in tracheal lumen in *Df(3L)BSC419* heterozygous (G) and homozygous (H) embryos is non-specific. Asterisk in

(A) indicates YFP signal from the *Dfd-GMR-YFP* balancer chromosome used to genotype embryos.

Scale bars (D-H): 50 µm.

**Figure S2 (related to Figure 1 and Figure 3) All M6 isoforms support TCJ formation.**

**(A–G)** *En face* view of lateral epidermis in control (*y w*; A) and in *M6*-deficient (*M6^MB02608^/Df(3L)BSC419*; B-G) embryos (stage 15) expressing UAS-M6-GFP (B), UAS- M6(B)-GFP (C), UAS-M6(C)-GFP (D), UAS-M6(D)-GFP (E), UAS-M6(E)-GFP (F), or UAS-M6(F)-GFP (G) in epidermal stripes under control of *en*-Gal4. Embryos were fixed and immunostained against Aka, M6, and Gli. Note that all M6 isoforms rescue TCJ localization of Aka and Gli. M6 Isoforms E and F show low expression levels and rescue Aka localization only partially.

Scale bar (A-G), 10µm.

**Figure S3 (related to Figure 4)**

**Palmitoylation does not significantly alter maintenance or mobility of M6 at TCJs.**

**(A)** Fluorescence recovery after photobleaching (FRAP) experiments in embryos (stage 15) homozygous for endogenous *GFP::M6* (control; top) or palmitoylation-deficient *GFP::M6*^Δ*Palm*^ (bottom). Time (min:sec) is indicated.

**(B)** Kymographs of series shown in (A) and quantification of fluorescence recovery indicate similar mobility of GFP::M6 (control; blue) and GFP::M6^ΔPalm^ (red).

**(C)** FRAP experiments in embryos (stage 15) expressing endogenous Aka::GFP in control (top) or in palmitoylation-deficient *M6*^Δ*Palm*^ embryos (bottom). Time (min:sec) is indicated.

**(D)** Kymographs of series shown in (C) and quantification of fluorescence recovery indicate similar mobility of Aka::GFP in control (blue) and in *M6*^Δ*Palm*^ (red) embryos.

Scale bars (A,C), 2 µm.

**Figure S4 (related to Figure 2)**

**Predicted orientation of M6 protein in the plasma membrane.**

**(A)** Structure prediction of M6-D from AlphaFold (ID: AF-Q9NGC6-F1) was analyzed using the PPM3 webservice to determine orientation of the protein in the membrane. The protein acquires a coiled-coil structure and is tilted 31 degrees with respect to the membrane perpendicular line (red dashed line).

**(B)** Same as in (A) but rotated by 180°. The three conserved juxtamembrane cysteines near the N-terminus are highlighted in red. A proline at position 13 (green) induces a kink that bends the N-terminus towards the plasma membrane.

**(C)** Close-up view of the palmitoylated region. Cysteines are shown in red, proline is shown in green. Arg11 and Thr85 from the second TMD engage in a hydrogen bond. Charged or polar side chains are highlighted.

## Supplemental Movies

**Video S1 (related to Figure 4)**

Dorso-lateral epidermis of stage 13 embryos homozygous for endogenous GFP::M6 (left) or GFP::M6^ΔPalm^ (right). Movies were aligned to the timepoint (t=30 min) that shows first enrichment of GFP signals at a minimum of 5% of the final number of vertices. Note the delay in vertex accumulation of GFP::M6^ΔPalm^ compared to the control. The movie of the control embryo ends at t=100 min. Time is indicated.

Scale bar: 50µm.

