## Supplementary figures and images for "Palmitoylation of proteolipid protein M6 promotes tricellular junction assembly in epithelia of *Drosophila*"

### Supplemental Figure 1

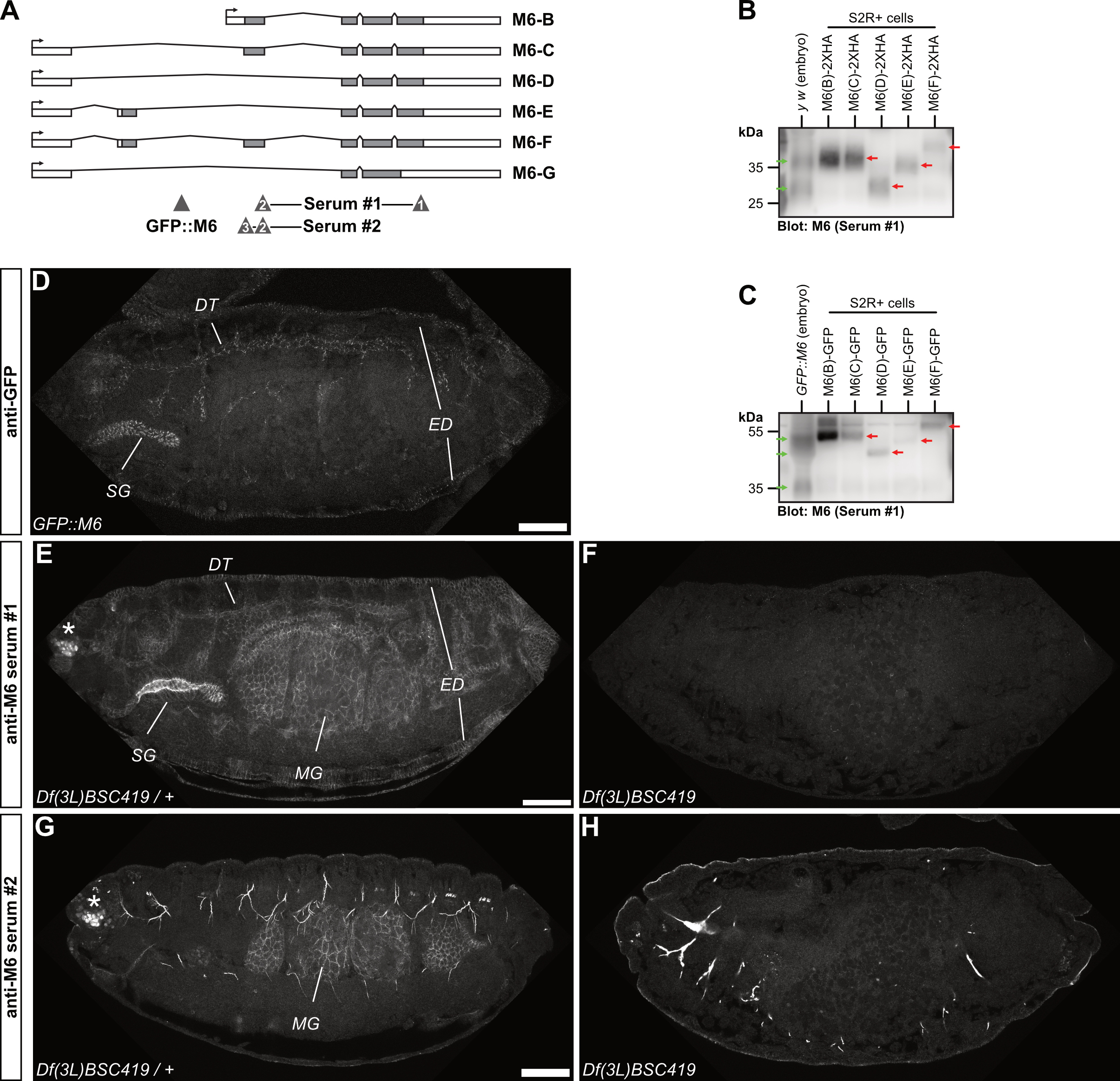

### Supplemental Figure 2

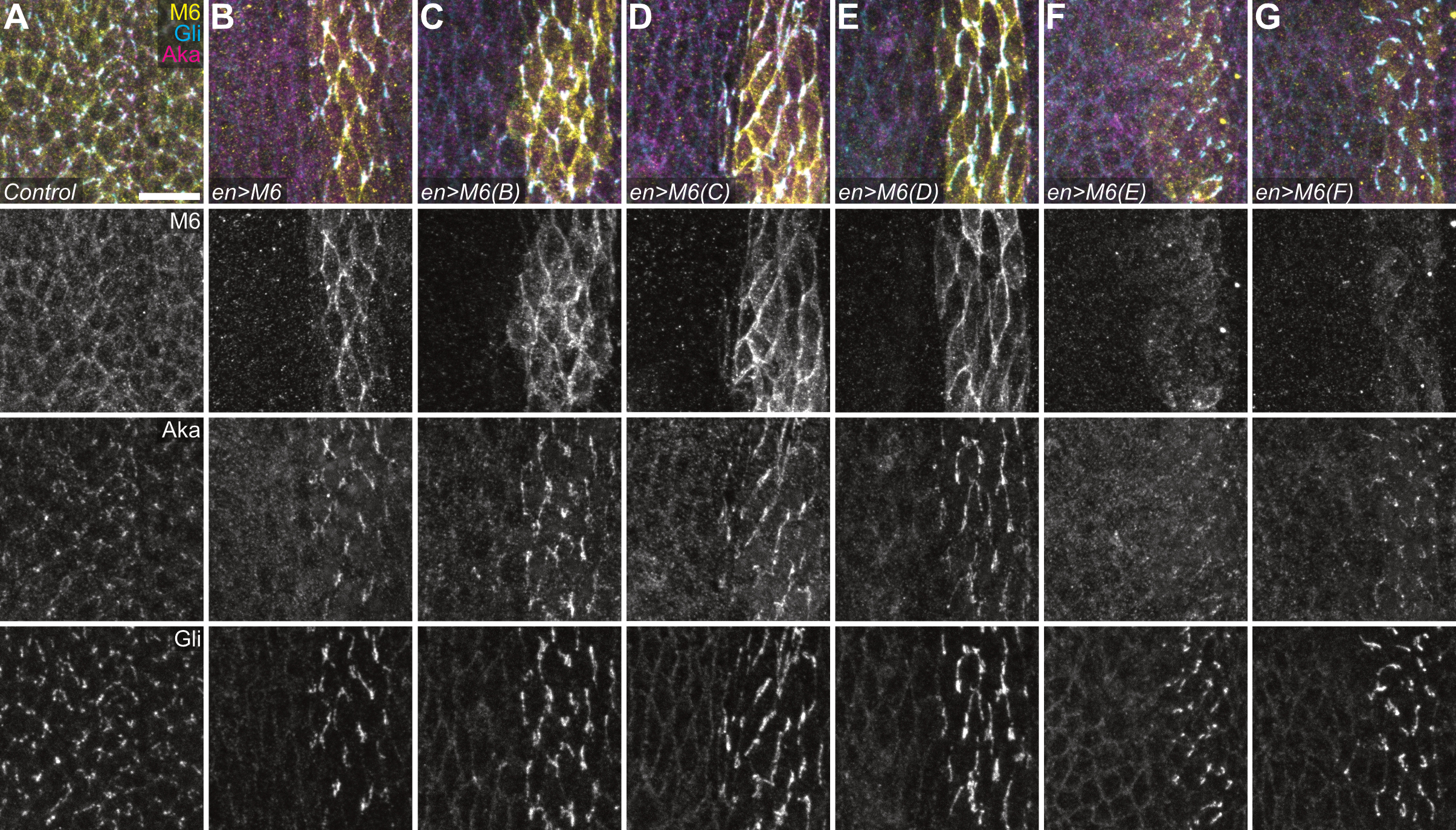

### Supplemental Figure 3

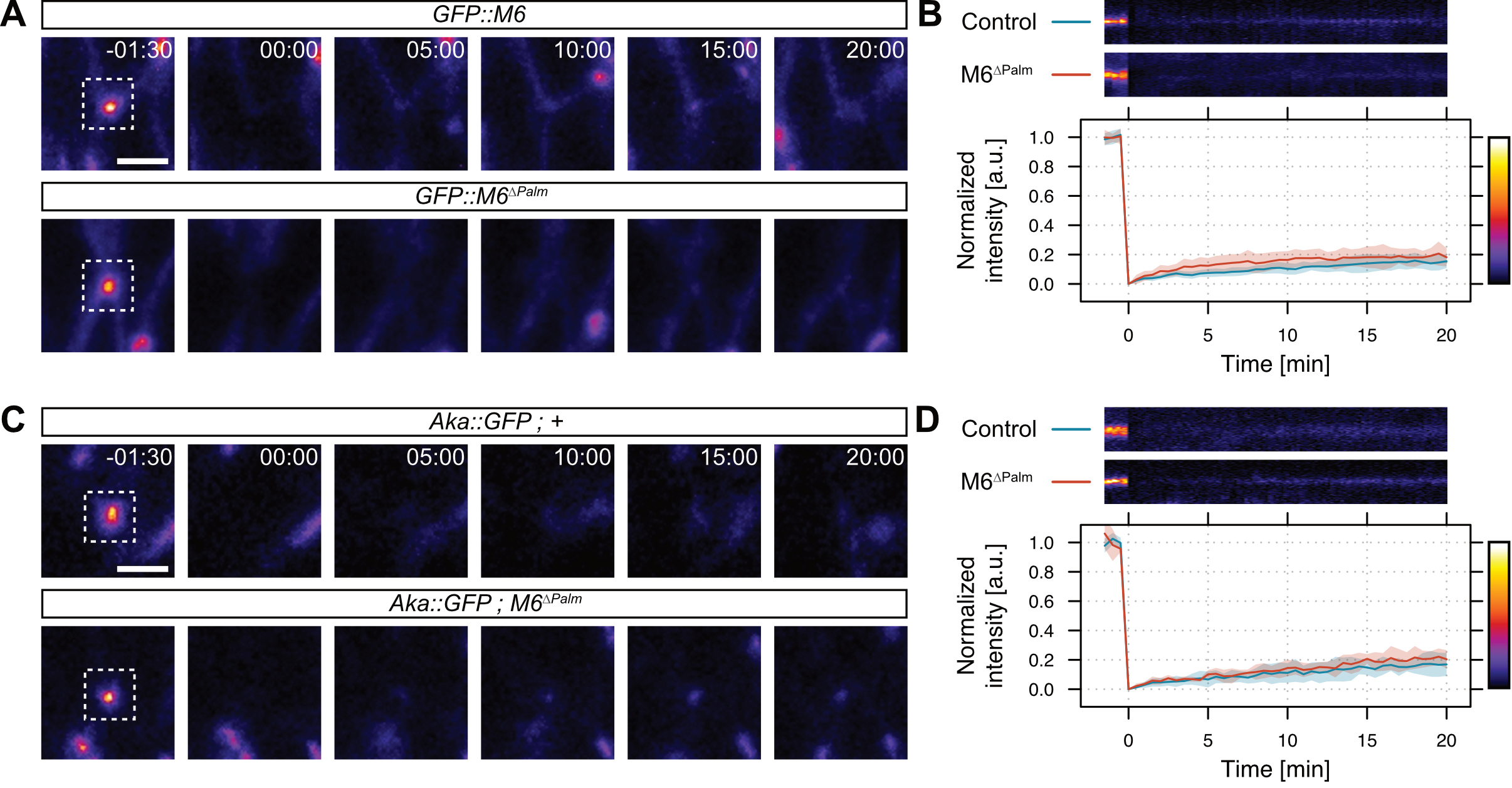

### Supplemental Figure 4

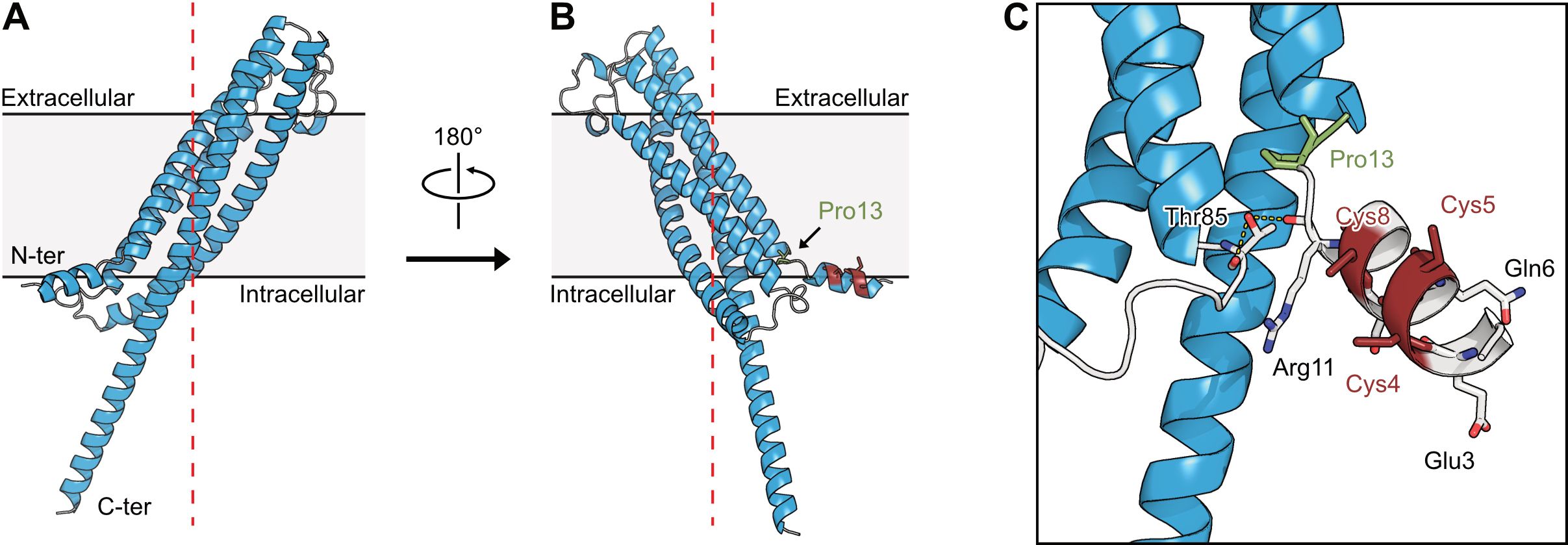
